# How motivational interviewing shifts food choices and craving-related brain responses to healthier options

**DOI:** 10.1101/2023.10.13.562241

**Authors:** Belina Rodrigues, Iraj Khalid, Solene Frileux, Benjamin Flament, Zeynep Yoldas, Martine Rampanana, Hippolyte Aubertin, Jean-Michel Oppert, Christine Poitou, Jean-Yves Rotge, Philippe Fossati, Leonie Koban, Liane Schmidt

**Author notes:** **Corresponding authors:** Belina Rodrigues and Liane Schmidt **Email:**. equal contribution (Solene Frileux, Benjamin Flament, Zeynep Yoldas, Martine Rampanana). **Author Contributions:** BR and LS conceived the original idea of the study and designed the study protocol and materials with advice from PF. BR, JYR, SF, IK, BF, JMO and CP included participants. BR collected data with help from MR, BF, SF, ZY, and IK. BR and HA analyzed the data under the supervision of LS and advice from LK. BR and LS drafted the manuscript, and PF and LK contributed to the final text. **Competing Interest Statement:** The authors declare no competing interest.

## Abstract

Changing one’s habits is challenging. Motivational interviewing (MI) has been proposed as a communication-based approach to overcome this challenge. Here, we tested how change and sustain talk suggestions, elicited by MI, influenced value-based dietary decision-making and responses of a recently developed neurobiological craving signature (NCS) in female adults. We found that following change talk, compared to sustain talk, participants’ food choices and activity in the ventromedial prefrontal cortex were more driven by the healthiness and less by the tastiness of food. These findings were paralleled by lower NCS responses to tasty food after change compared to sustain talk. Further, following change talk, participants’ body mass indices moderated the NCS decoding of healthy and tasty food choices. These results show that MI can shift value-based decision-making and reduce craving-related brain marker responses to highly palatable food items. The findings contribute to a better understanding of behavioral change interventions toward healthier eating.

## Introduction

Modifiable behaviors such as tobacco use, physical inactivity, unhealthy eating habits, and harmful alcohol consumption increase the risk of non-communicable diseases and account for a significant percentage of worldwide disability and mortality (1–3). In particular, unhealthy eating habits are associated with an increased risk of all-cause mortality (4) and carry a significant economic burden greater than smoking and physical inactivity (5). However, unhealthy eating behavior is rising worldwide (6).

Given the consequences of unhealthy dietary habits, it is essential to understand the mechanisms of interventions designed to promote behavior change. In this study, we investigated the neurocognitive effects of a prominent communication-based approach to behavior change – motivational interviewing (MI) (7–11). MI is a collaborative clinical method, a goal-oriented style of communication focusing on the language of change. It aims to strengthen motivation and commitment to a specific goal. In more detail, MI elicits personal statements for change, called change talk, and deals with reasons against change, i.e., sustain talk, within an atmosphere of acceptance and compassion (12). Research on the associations between MI and behavioral outcomes showed that a higher proportion of change talk over sustain talk is associated with reduced risk-seeking behavior (13, 14).

Previous work on the neural correlates of MI has focused on substance use disorders such as alcohol and cannabis use. These studies have provided convergent evidence that listening to change talk increased the recruitment of brain regions associated with cognitive regulation in terms of self-control among cannabis users (15) and dampened the activation of brain areas related to craving and reward when viewing alcohol-related cues (16). These findings suggest that MI might generate a cognitive shift towards self-control and a reduction in craving. However, there is a lack of consensus about the joint behavioral and neural effects when participants mentally transition between change and sustain talk during decision-making. Further, no previous studies have investigated whether MI can alter responses in *a priori* and independently established brain markers of craving, such as the recently developed neurobiological craving signature (17).

Self-control propensities have been shown to play a crucial role in advantageous economic choices (18). Models of economic choice propose that the decision process involves two stages (19). During valuation, various attributes of choice alternatives are integrated into stimulus values, which approximate a decision-maker’s preferences. Stimulus values are then compared during action selection to lead to a preferred choice. Findings from research on the neurobiology of valuation show that value is computed within the brain’s reward and valuation system formed by the vmPFC, ventral striatum, and posterior cingulate cortex (for review, see (20, 21)). These value computations can be modulated by self-control, defined by the ability to forgo tasty and liked food in favor of healthier food to improve the quality of eating habits. Such value modulation has been reported to involve the activation of the dlPFC (22).

However, sticking to more self-controlled, healthier eating is tricky in everyday life. The vmPFC has also been shown to encode sensitivity to food stimuli and food craving (23–25) and has strong positive weights in a recently validated neural signature of food and drug craving – the NCS (17). Food cravings relate to eating and weight gain over time (26). Motivational interviewing has been found effective for such situations with prominent cravings, for example, substance use disorders (11). However, its effectiveness to favour the self-control of dietary decision-making and/or reduce neural decoding of food craving is unknown.

Here, we adopted this value-based decision-making framework and used the neurobiological craving signature (NCS) to predict that listening to change versus sustain talk should (1) change food valuation during dietary decision-making by shifting the weights of healthiness and tastiness of food, (2) alter how the vmPFC as a central hub of the brain’s valuation system encodes these attributes of food during economic choice formation, and (3) reduce craving related brain responses as quantified by the NCS to tasty food items.

## Results

### Behavioral results

We first tested whether change compared to sustain talk affected the value assigned to food stimuli, i.e., how much participants wanted to eat the food items. Two analogous linear mixed effects models (LME) of stimulus value were used: LME 1 was fitted to stimulus values assigned to healthy and tasty food. This model revealed that change talk significantly increased food value compared to sustain talk (main effect of change versus sustain talk: β = 0.05, SE = 0.02, 95%CI [0.006 – 0.08], t = 2.3, p =0.02). The model further showed a non-significant main effect of type of attribute (healthiness versus tastiness: β = -0.03, SE = 0.03, 95% CI [-0.09 – 0.027], t = -1.1, p = 0.29) and a significant interaction type of talk by attribute type (β = 0.06, SE = 0.02, 95% CI [0.03 – 0.09], t = 3.64, p =0.0004) such that after change talk (compared to sustain talk), stimulus value was greater for healthy food items. LME 2 was fitted to stimulus values assigned to unhealthy and untasty food. This model also found a significant interaction talk by attribute type (β = -0.06, SE = 0.02, 95% CI [-0.11 – -0.01], t = -2.5, p = 0.01), but a non-significant main effect of talk (β = -0.04, SE = 0.03, 95% CI [-0.09 – 0.012], t = -1.5, p = 0.14), and a significant main effect of attribute type (β = 0.3, SE = 0.03, 95% CI [0.20 – 0.34], t = 7.9, p = 1.3e-12). Both models were controlled for BMI.

To describe these interactions in more detail, posthoc t-tests and Figures 1a and 1b show that healthy food was chosen more often (t(31)= 4.1, p <.001, paired, two-tailed t-test, Cohen’s d = 0.72), and unhealthy food was rejected more often (t(31) = -2.4, p = 0.02, paired, two-tailed t-test, Cohen’s d = -0.42) after change talk than after sustain talk. This difference in stimulus values was non-significant for tasty foods (change versus sustain talk) (t(31) = -0.6, p = 0.55, paired, two-tailed t-test, Cohen’s d = -0.1) nor for untasty foods (t(31)= 0.81, p = 0.42, paired, two-tailed t-test, Cohen’s d = 0.14).

**Figure 1.**
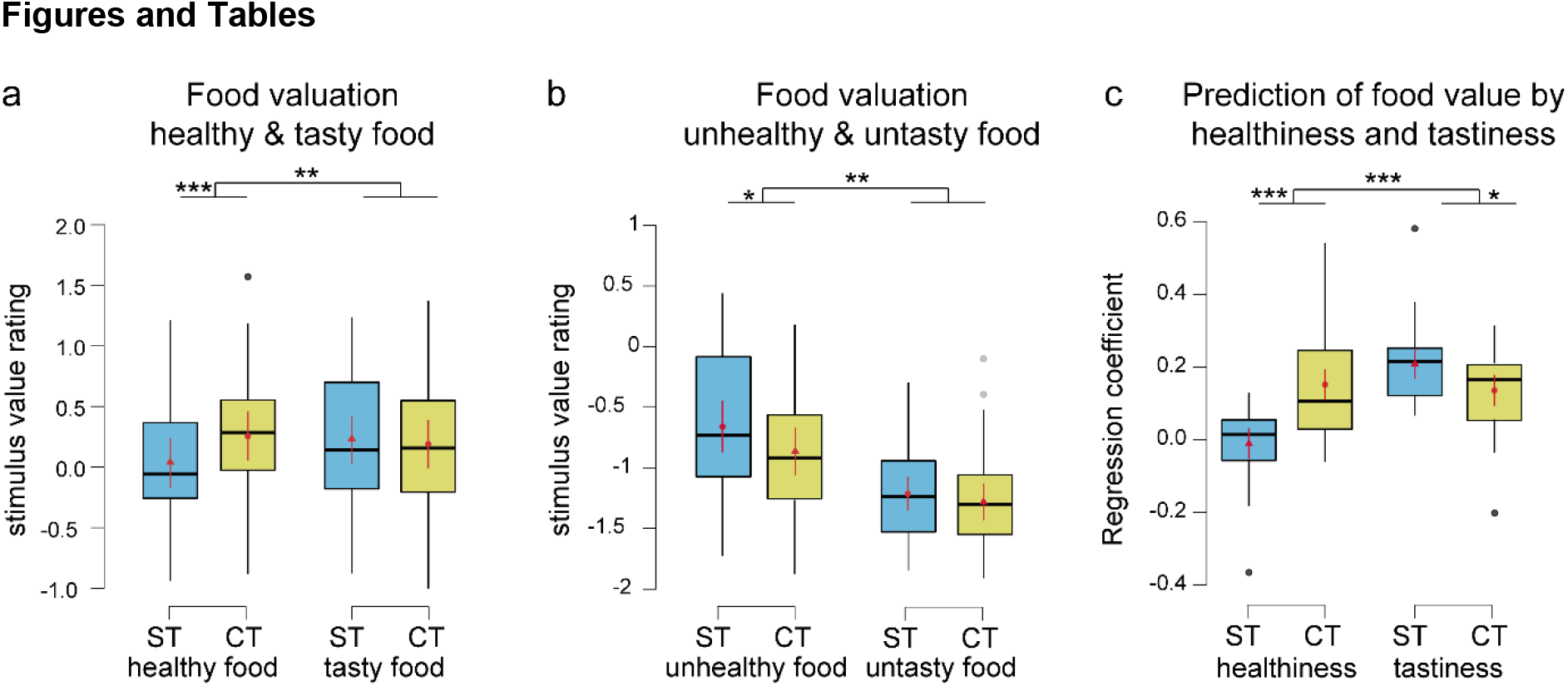
Behavioral results in N=32 participants. Boxplots display 95% CI for value assigned after change (CT) and sustain (ST) talk to **(a)** food stimuli during healthy and tasty food choices, **(b)** value assigned to food stimuli during unhealthy and untasty food choices, and **(c)** the prediction of food stimulus value by tastiness and healthiness ratings across all food choices. Change talk (and compared to sustain talk) increased the weight of healthiness on the food choice which translated into choosing more often healthy and rejecting more often unhealthy food. ^***^ p<0.001, ^**^p<0.01, ^*^ p<0.05 for a significant interaction attribute type (healthiness versus tastiness) and talk type (change versus sustain talk) and paired, two-tailed t-tests.

To further test how listening to change versus sustain talk influenced how much healthiness and tastiness information were integrated into stimulus values across all food choice trials, the individual beta values from a general linear model fit to stimulus value ratings were entered into a second-level analysis of variance (*SI Appendix* Table S1). Specifically, the ANOVA tested how much first-level interaction terms of interest (i.e., CTxTR, CTxHR, STxTR, and STxHR) differed as a function of the type of talk (change versus sustain talk) and type of attribute (healthiness versus tastiness).

In line with food valuation-related analyses, the second-level random effect ANOVA revealed a main effect of attribute type (F(1,124)=22.6, p<0.001, *η*^2^ = 0.14), a small borderline effect of talk (F(1,124)= 4,12, p=0.045, *η*^2^ = 0.0^2^), and a significant interaction talk by attribute (F(1,124) =32.6, p<0.001, *η*^2^ = 0.15) (Figure 1c). As shown in Figure 1c, after listening to a change talk, healthiness predicted significantly more food stimulus value than after listening to a sustain talk (CTxHR vs STxHR: t(31) = 3.9, p<0.001, Cohen’s d = 0.7, paired, two-tailed t-test). In contrast, after listening to a sustain talk, the tastiness predicted more food stimulus values than after listening to a change talk (STxTR vs CTxTR: t(31)= 2.3, p=0.029, Cohen’s d = 0.4, paired, two-tailed t-test).

In summary, these complementary statistical models converged on the finding that change talk compared to sustain talk increased the participant’s propensity to consider the healthiness more and the tastiness less during food valuation.

### Effects of talk on food value computation within the prefrontal cortex

At the time of food choice, significant activation within the brain’s reward and valuation system encompassing the vmPFC, ventral striatum, and posterior cingulate cortex correlated significantly with food stimulus value (*p*_FWE_ < 0.05 family-wise error correction on the peak and cluster level, Figure 2a, *SI Appendix* Table S2). This activation did not significantly differ between change and sustain talk conditions. We next tested whether listening to change versus sustain talk affected the integration of healthiness and tastiness within the value-encoding vmPFC region of interest. As shown in Figure 2b, the vmPFC-value encoding region of interest correlated stronger with healthiness at the time of food choice after listening to change compared to sustain talk (small volume correction, *p*_FWE_ < 0.05). The vmPFC ROI correlation to tastiness did not differ between change and sustain talk conditions.

**Figure 2.**
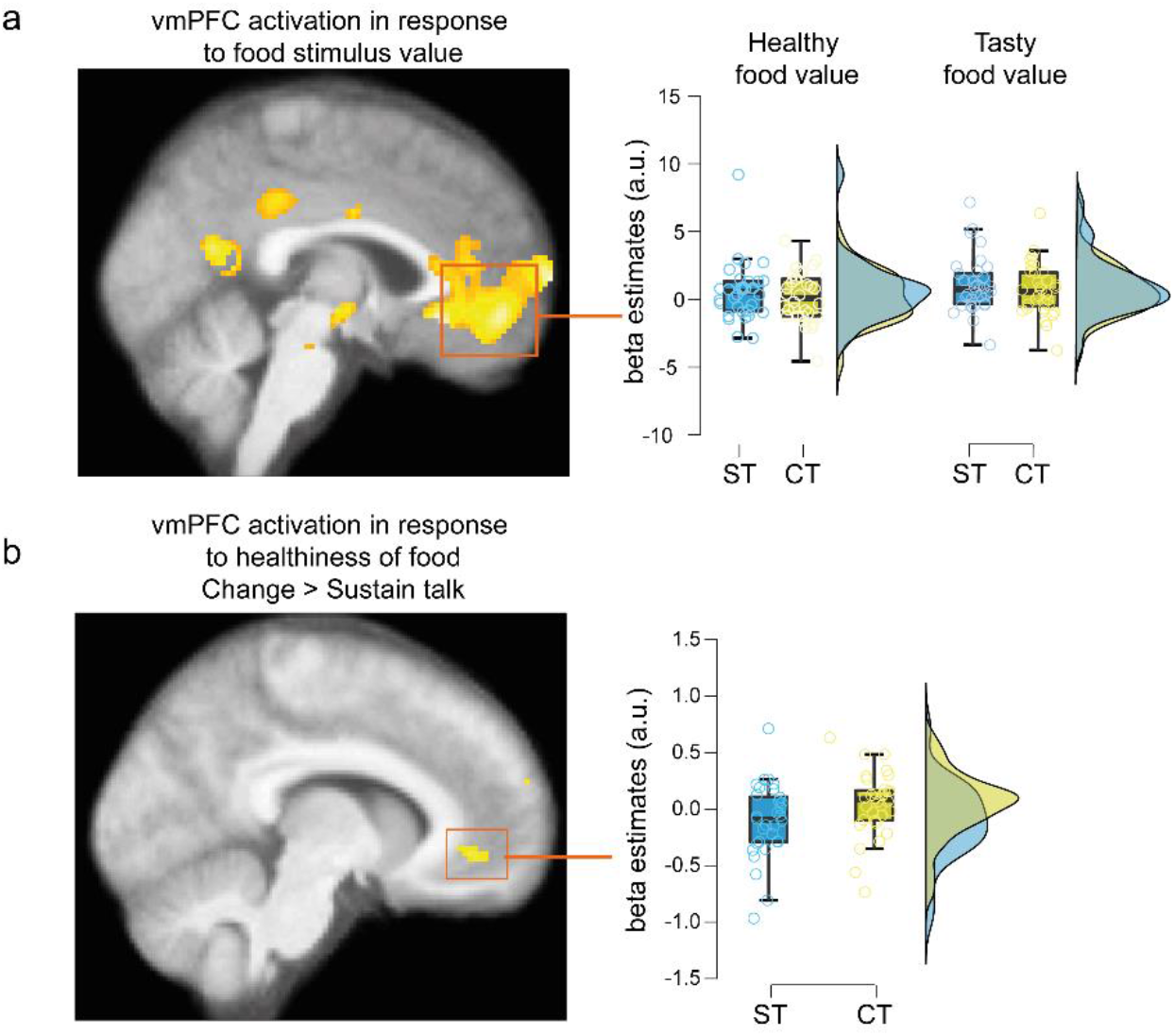
Brain responses to stimulus value and healthiness. **(a) Stimulus value encoding in the brain**. Statistical parametric maps show brain activation in correlation to stimulus value at the time of food choice. Significant voxels in yellow and orange survived a whole brain threshold of p_FWE_ < 0.05, family-wise error corrected on the peak level, and are shown for display at p<0.001 uncorrected. **(b) Contrast between change versus sustain talk for healthiness encoding in the brain**. SPMs show activation of the vmPFC region of interest that correlated stronger to healthiness ratings at the time of food choice after the change than after sustain talk. Significant voxels in yellow survived small volume correction (SVC p_FWE_ < 0.05) within the vmPFC and are displayed at an uncorrected whole brain threshold of p<0.005. The boxplot shows 95% confidence intervals of beta values extracted from the peak activation in response to healthiness ratings within the vmPFC (MNI_peak_ =-8, 46, -8) for illustration. Colored dots correspond to individual participants. All SPMs are superimposed on the average structural brain image.

### Prediction of food stimulus value by the neural craving signature (NCS)

We next tested whether the observed behavioral effects of change and sustain talk on healthy and tasty food choices were paralleled by differential responses to food stimulus values (i.e., choices) of the recently validated neurobiological craving signature (17).

A linear mixed effects model (LME 3), analogous to LMEs 1 and 2, was fit to the NCS responses to food stimulus value. The model revealed that overall NCS responses were higher for tasty food choices than healthy food choices (β = -1.9, SE = 0.63, 95% CI [-3.2 – -0.69], t = -3.1, p = 0.0025). The model also indicated a non-significant effect of talk (β = -0.07, SE = 0.32, 95% CI [-0.70 – 0.56], t = -0.21, p = 0.83) and, in line with our prediction, a significant interaction talk by attribute (β = 0.86, SE = 0.32, 95% CI [0.23 – 1.5], t = 2.7, p = 0.008). This interaction effect (see Figure 3b) was driven by a significant reduction in NCS responses to tasty food choices in the change, compared to the sustain talk condition (t(31) = -1.9, p = 0.03, one-tailed t-test, Cohen’s d = -0.35). In addition, the NCS tended to respond more to healthy food choices following change talk than following a sustain talk (t(31) = 1.5, p = 0.07, one-tailed t-test, Cohen’s d = 0.28).

**Figure 3.**
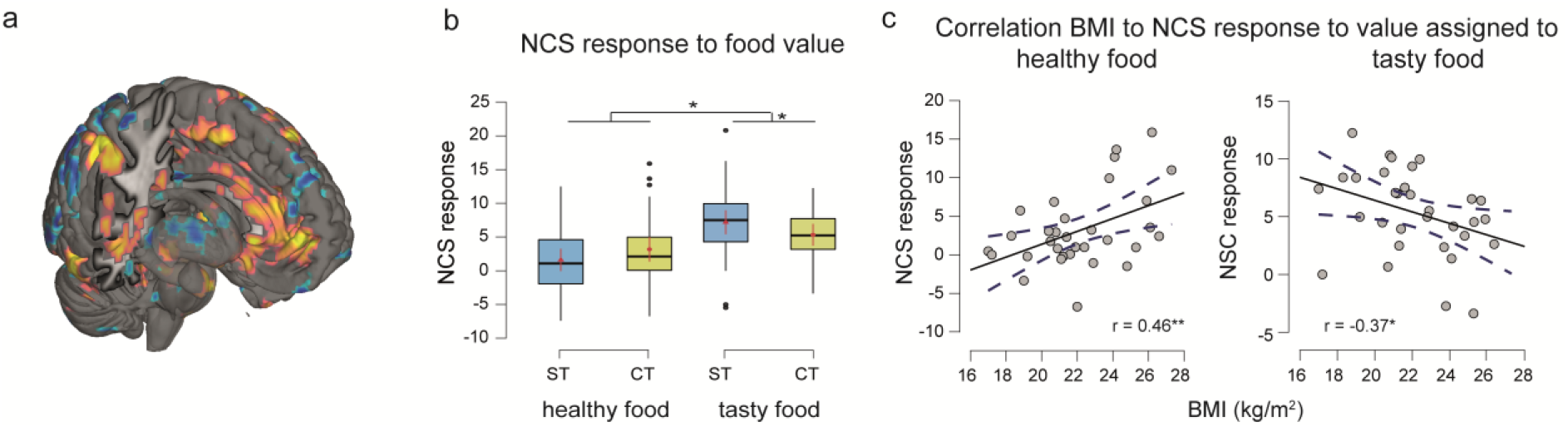
Neural craving signature prediction of food stimulus value in change and sustain talk conditions. **(a)** Pattern of the neural brain signature of craving (NCS) with positive weights displayed in yellow and red and negative weights shown in blue (at p<0.05 uncorrected for display). **(b)** The boxplots depict 95% CI for the NCS response to food stimulus value during healthy and tasty food choice trials. Error bars correspond to standard errors. Horizontal bars indicate the median, and red points mean values. Individual points in black indicate outlier participants. **(c)** Correlations between BMI and the neurobiological craving signature (NCS) response to the value assigned to healthy and tasty food in the change talk condition.

Lastly, we explored the role of inter-individual differences in NCS responses to healthy and tasty food choices after change and sustain talk. In the change (but not in the sustain) talk condition, body mass index (BMI) correlated significantly and positively with the NCS responses to healthy food choices (Pearson’s r = 0.46, p = 0.009). On the contrary, it correlated negatively with NCS responses to tasty food choices (r = -0.36, p = 0.04). This suggests that the effects of change talk on the NCS might be especially pronounced in participants with higher body mass.

## Discussion

Motivational interviewing is an evidence-based intervention technique in the context of addiction and behavioral change more broadly. This study aimed to characterize the effects of motivational interviewing on dietary decision-making and craving-related brain responses to food choices. We combined fMRI with behavioral testing of value-based dietary decision-making to contrast in a within-participant design two types of suggestions, i.e., one in favor, the other against the change of eating habits elicited by motivational interviewing.

In line with our predictions, change talk led to a greater weighting of the healthiness of food items and a lower weighting of tastiness, an effect that was paralleled by predefined craving-related responses in the neural craving signature (NCS) – a recently established brain marker of food and drug craving (17). Moreover, the participant’s body mass index moderated the NCS responses to tasty and healthy food choices. These findings suggest an interaction between motivational interviewing and body weight on craving-related brain circuits.

Previous research has looked at the neural mechanisms of motivational interviewing in the context of alcohol and substance use (see (27–29) for examples). Here, we investigated by using univariate and multivariate brain-behavior analyses an aspect that has yet to be reported: food preferences, which were measured by an incentive-compatible value-based decision-making task. Sticking to healthier food choices is difficult, and eating-related conditions such as diabetes are a worldwide problem. Here, we tested in hungry, normal to overweight participants the cognitive effects of shifting between change and sustain talk in the laboratory context. Our findings in normal to overweight to participants are therefore a proof of concept for motivational interviewing to affect the basic mechanisms of value-based dietary decision-making. They further align with an increased focus on prevention and the assumption that motivational interviewing might also be helpful as a prevention strategy instead of an adjunct treatment strategy (30). Future work should test how the cognitive shifting between change and sustain talk modulates food valuation and valuation-related brain activation in populations struggling with food-addiction-like behavioral habits and obesity.

Our behavioral finding of value modulation suggests that change talk might have generated a form of dietary self-control (22, 31–36). Previous studies of dietary decision-making and its self-control have used instructions to incite participants to shift explicitly between strategies, such as distancing from or focusing attention on the healthiness or tastiness of food during incentive compatible dietary decision-making tasks (22, 31–36). In line with this past work participants of the current study performed a similar incentive compatible task in the fasted, hungry state, which may have favored more indulgent decision-making under sustain talk conditions, and at the same time may have also enhanced the conflict between short-term taste and long-term health rewards of food under change talk conditions. These two aspects are important ingredients to measure self-control propensities during dietary decision-making. However, the current study also differed from this past work in several, important and novel aspects. Change and sustain talk was not always related to health or taste but to other aspects of food behavior (e.g., environmentally friendly food behavior). These reasons were unique and personal to each participant. Moreover, participants were told to make food choices following their natural preferences and to remember the suggestions they listened to. The instructions also emphasized that both types of suggestions, change and sustain talk, were valid and that they should not favor one type of talk over another. Our findings thus suggest that patients and health professionals do not need to focus on the “health aspect” to elicit behavior change but instead on the personal reasons most relevant to a patient. This is not trivial, as not all patients are health-focused. While it remains an open question to what extent the observed behavioral effects generalize to real-world food choices, the scope of our study was to shed light on the primary cognitive and neural effects of shifting between change and sustain talk during economic food choice in a controlled laboratory setting.

The behavioral results were underpinned by the univariate finding that the vmPFC encoded food value during dietary decision-making irrespective of the type of talk. This result is in line with prior evidence that the vmPFC is a hub within the brain’s valuation system that encodes expected and experienced rewards across different economic choice problems (37–39). On the behavioral level, the type of talk (i.e., change versus sustain talk) influenced how food preferences were formed, which was approximated by the integration of tastiness and healthiness into food stimulus values (i.e., food choices). In line with this behavioral finding, the value-encoding vmPFC region of interest correlated stronger to the healthiness of food at the time of choice following change talk than following sustain talk. This result aligns with findings from neuroeconomics that showed that the vmPFC computes the value signals that drive economic choices (20). Past work from this stream of research has also found another brain region, the dorsolateral prefrontal cortex (dlPFC), to play a role in value modulation induced by cognitive regulation such as self-control during dietary decision-making (31, 32, 36). Here, the univariate brain analysis results were restricted to the vmPFC. We did not find any evidence for the contribution of the dlPFC to dietary decision-making after change and after sustain talk. Different experimental designs, participant samples, and contextual conditions might explain these odds. For example, the dlPFC might be more prominently recruited during economic choice formation when participants face a real-world conflict between change and sustain talk such as struggling with food addiction or obesity. Alternatively, the dlPFC might also be more implicated in contexts where instructions to focus on specific attributes of food are more explicit.

Alternatively, our results that change relative to sustain talk affects the integration of healthiness and tastiness into stimulus value might be better explained by the recruitment of craving-related activation in more than one or two brain regions (40). Current multivariate, statistical approaches in the analysis of fMRI data use machine-learning algorithms to build predictive models – brain signatures to decode behavior from whole brain activation (40). Here, we leveraged this approach and used the neurobiological craving signature (NCS). The NCS encompassed value and reward-related brain regions such as the vmPFC and ventral striatum, but also other regions such as the insula, anterior midcingulate cortex, and temporal/parietal association areas (17). It has been cross-validated to predict individual food and drug craving levels across different participant samples of drug and non-drug users to whom both drug and high palatable (i.e., tasty) food cues were presented. We showed that NCS predictions of healthy and tasty food choices were different following change compared to sustain talk. In more detail, similar to the effects of talk type on stimulus value, we found a significant interaction type of talk by attribute on the NCS response. NCS responses to *tasty* food choices were significantly reduced in the change compared to the sustain talk condition. This finding complements the effects of talk on stimulus value ratings, which was driven by a significant difference between change and sustain talk on *healthy* food choices. These differences might be explained by the fact that the NCS was trained to predict cravings for highly palatable food items and might, therefore, be more sensitive to tasty food choice trials in our data.

Moreover, the NCS responses to healthy and tasty food choices after change talk varied as a function of BMI. While these correlations should be interpreted with caution given the small sample size, they suggest that participants with higher BMI present greater NCS responses to healthy food choices and lower NCS responses to tasty food after change talk. Brain activation, especially within the brain’s value and reward systems, which are a prominent part of the NCS, is influenced by weight status and hormones that signal energy states, such as leptin, which are directly related to body fat (41–43). Therefore, links between the brain, body composition, and energy-store signaling hormones could explain our correlational findings. Change talk could make craving- and reward-related brain circuits more sensitive to such internal, afferent signals from the periphery. However, while these correlations were non-significant for sustain talk conditions, this might be due to the small sample size. Thus, these correlational results should be replicated in larger sample sizes that also include participants with obesity.

A few limitations need to be considered. Here, we report on the influence of client language on dietary decision-making. However, motivational interviewing is more than just eliciting change talk. It comprises two sets of skills: technical and relational (44). The technical skills, composed of open questions and reflections, elicit and strengthen change talk. The relational skills, comprising aspects such as empathy, ensure an atmosphere of acceptance and compassion. Both features aim at increasing the frequency and strength of change talk while handling sustain talk reasons (44, 45). Interestingly, Magill et al. (2018) found a non-independence of technical and relational components of motivational interviewing, as the variance of the technical element was partly explained by varying levels of empathy. It has also been found that the relational aspect predicted more reflections of change, which, in turn, predicted more change talk (46). These findings point to a synergistic effect of both sets of skills. However, the current study design combining value-based decision-making with fMRI did not allow for assessing the impact of change talk in conjunction with these relational components on real-world behavioral change propensities.

The scope of our findings is restricted to the basic cognitive mechanisms of decision-making. In the real world, the medical appointment between patient and health professionals and the post-session processes may elicit many effects (16), which drive behavior change. Additionally, future studies should replicate our findings in other samples. Moreover, other behavior change approaches should be contrasted to motivational interviewing effects by future work to determine the specificity of basic cognitive mechanisms and how they might give rise to real-world behavioral change in the long term (47).

In conclusion, our findings indicate that MI leads to a shift in dietary decision-making paralleled by changes in specific brain circuits related to value-based decision-making and craving. These findings advance our understanding of the neurocognitive mechanism of communication-based approaches to behavior change, which may have a place in regular medical appointments to facilitate sticking to healthier eating habits.

## Materials and Methods

### Ethical considerations

The study was approved by the local ethics committee and conducted in accordance with the Declaration of Helsinki. All participants provided written and informed consent.

### Study-pre-registration

This study was part of a still ongoing, larger-scale pre-registered study (https://clinicaltrials.gov/study/NCT05101863). It focused exclusively on the participants whose BMI was less than 30. A second deviation from the pre-registered protocol involved using a more recently validated neurobiological craving signature (17).

### Participants

A total of 40 participants were recruited via public advertisement. The inclusion criteria were to self-identify as female, aged between 18 and 70 years old, right-handed, with normal to corrected-to-normal vision, no history of substance abuse or neurological or psychiatric disorder, no medication, and no MRI contraindication. Inclusion was limited to self-identified female participants to have a more homogenous group. Research on the physiology of appetite and eating behavior has shown that men and women differ in their food behavior (48–52), such that women have been shown to display stronger brain activation to appetite-enhancing food stimuli (48, 51), despite a strong propensity for dietary self-restraint (51).

Eight participants were excluded from analysis due to problems with the MRI scanner (n=5), computer software to record the behavioral data (n=2), and the discovery of an fMRI contraindication on the day of the MRI visit (n=1) (Table 1).

**Table 1.**
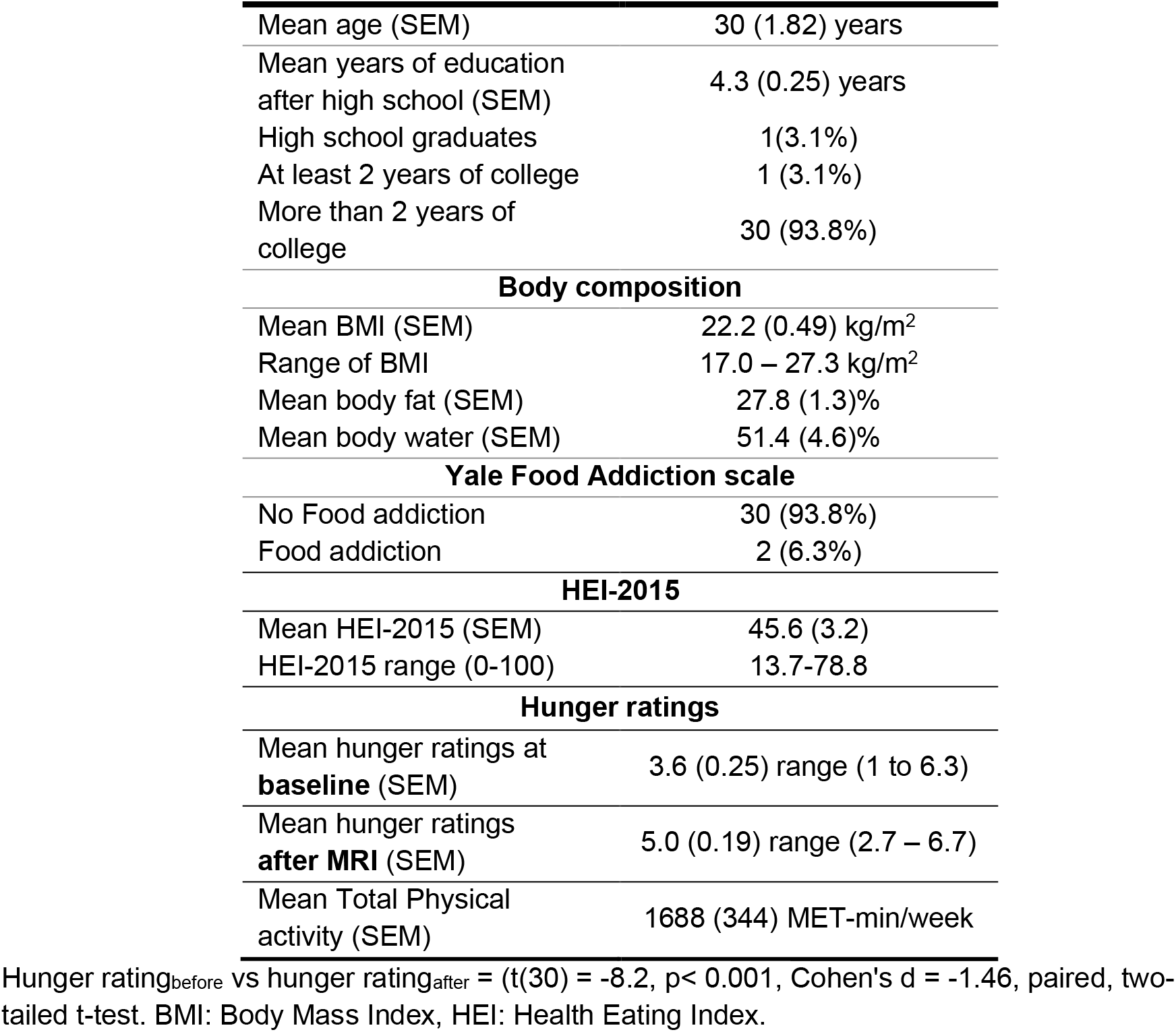
Socio-demographics and body composition of n=32 participants.

|  |  |
| --- | --- |
| Mean age (SEM) | 30 (1.82) years |
| Mean years of education after high school (SEM) | 4.3 (0.25) years |
| High school graduates | 1(3.1%) |
| At least 2 years of college | 1 (3.1%) |
| More than 2 years of college | 30 (93.8%) |
| **Body composition** |  |
| Mean BMI (SEM) | 22.2 (0.49) kg/m<sup>2</sup> |
| Range of BMI | 17.0 – 27.3 kg/m<sup>2</sup> |
| Mean body fat (SEM) | 27.8 (1.3)% |
| Mean body water (SEM) | 51.4 (4.6)% |
| **Yale Food Addiction scale** |  |
| No Food addiction | 30 (93.8%) |
| Food addiction | 2 (6.3%) |
| **HEI-2015** |  |
| Mean HEI-2015 (SEM) | 45.6 (3.2) |
| HEI-2015 range (0-100) | 13.7-78.8 |
| **Hunger ratings** |  |
| Mean hunger ratings at **baseline** (SEM) | 3.6 (0.25) range (1 to 6.3) |
| Mean hunger ratings **after MRI** (SEM) | 5.0 (0.19) range (2.7 – 6.7) |
| Mean Total Physical activity (SEM) | 1688 (344) MET-min/week |

Hunger rating<sub>before</sub> vs hunger rating<sub>after</sub> = (t(30) = -8.2, p< 0.001, Cohen's d = -1.46, paired, two-tailed t-test. BMI: Body Mass Index, HEI: Health Eating Index.

### Experimental procedure

The study was composed of two visits that took place one week apart.

#### Visit 1

Upon enrolment, sociodemographic data such as gender, age, and level and years of education and highest obtained diploma were collected. Then, each participant underwent a 24h dietary recall to study food intake according to the manual adaptation of the Automated Multiple-Pass Method (53). After obtaining a list of all food items eaten on the previous day, the interviewer probed for frequently forgotten foods such as savory snacks or candies. Then, the time and name of the eating occasion (e.g., breakfast) and a detailed description of the type of food item, cooking method, and other relevant aspects. Lastly, a final revision of the dietary intake took place (53). Whenever possible, visit 1 was scheduled so that seasonal changes (e.g., weekends, holidays such as Christmas) in the dietary habits would not be captured.

Following the 24h dietary recall, all participants underwent a semi-structured brief (i.e., 20-30 minutes) motivational interviewing session, which was divided into four phases: The interviewer, who was a dietitian trained in MI, elicited the participants to reflect on the following: (1) Their dietary patterns and whether there was a desire for a change, and (2) the advantages and disadvantages of the behavior they would like to change. The experimenter then asked participants to (3) rate the importance and their readiness for change on a 10-point Likert scale and 4) verified again whether they still would like to change an aspect of their dietary habits. The brief motivational intervention was audio-recorded, and 5 change and 5 sustain talk statements were extracted for use during an fMRI session at the second visit one week later (for examples see *SI Appendix* Table S3). Phrases were chosen by agreement of two different raters trained in MI.

#### Visit 2

A week after the first visit, participants underwent an fMRI session after overnight fasting. Before the fMRI session, and to ensure a balanced distribution of healthy, unhealthy, tasty, and untasty food stimuli across the two talk conditions, participants rated each food item in terms of healthiness and tastiness on a 4-point scale: strong no, no, yes, strong yes. Additionally, hunger ratings were measured on a 7-point Likert scale before and after the fMRI session. Importantly, these hunger ratings were averaged across the rating of three aspects of hunger: hedonic (i.e., How pleasant would it be to eat, now?), homeostatic (i.e., How much could you eat, now) and general (i.e., How hungry are you, now?).

To study the effect of change language, a modified version of a well-known value-based dietary decision-making task was presented to the participants (22, 31, 32, 35). It comprised 10 blocks of choice trials: Five blocks started with change talk statements and five with sustain talk statements. The order of change and sustain talk blocks was randomized. Each block counted 15 decision-making trials for 75 trials per talk condition. The blocks started by instructing the participants to keep in mind the talk statement they will listen to. Then, a 15-second talk statement was played, and the corresponding verbatim could be read on the screen. Participants were asked to read and listen to their voices expressing the change or sustain talk statements, respectively. After listening and reading the suggestions they rated 15 food stimuli of varying tastiness and healthiness on whether they wanted to eat this food for real at the end of the experiment. Each choice trial started with the display of a food stimulus, and participants had three seconds to rate whether they wanted to eat the food item for real at the end of the experiment by using a 4-point Likert scale ranging from strong no to strong yes. Trials were separated by a jittered fixation cross (Figure 4). A short break was planned after 75 trials (i.e., 5 blocks) to reduce head movement. A brief practice session occurred outside the MRI beforehand, and a second practice session occurred within the MRI to adjust the headphones’ volume.

**Figure 4.**
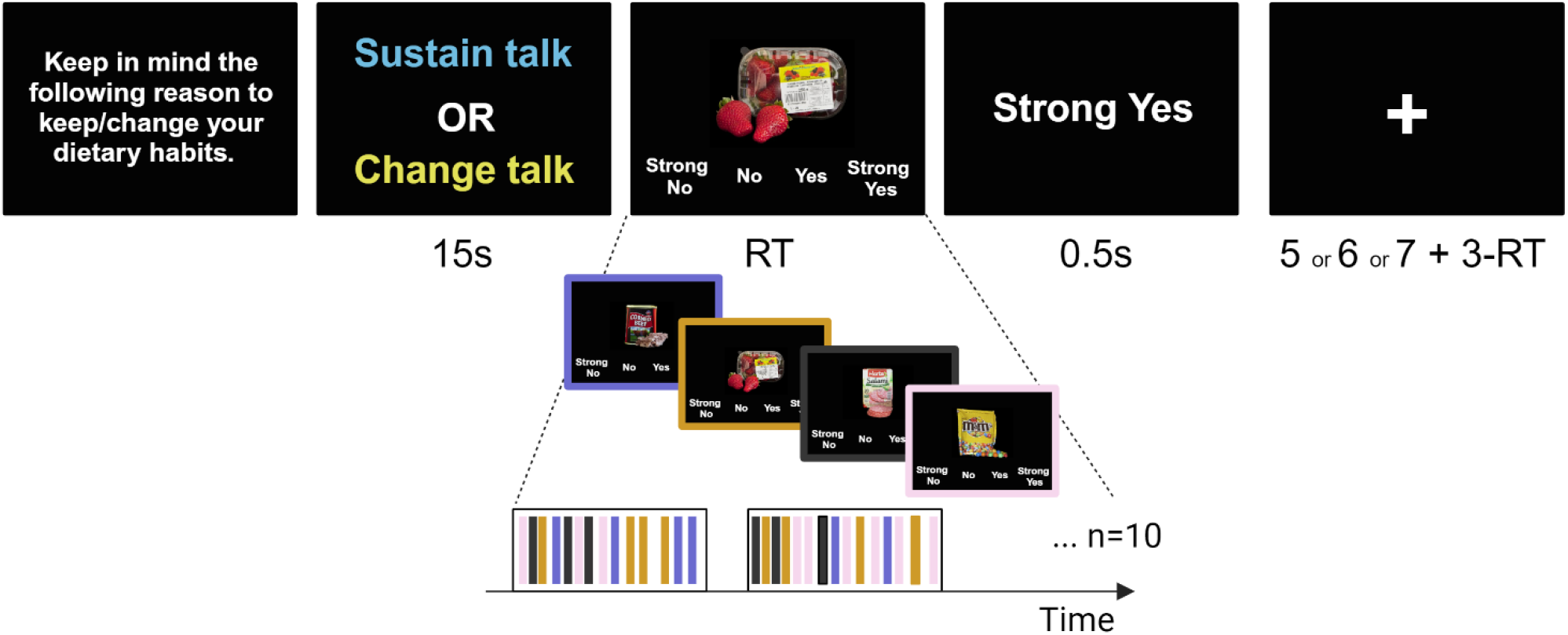
Dietary decision-making task. Panels display screenshots of successive events within a change and sustain talk blocks. Each block started with a reminder to keep in mind the statement related to change/sustain current dietary habits. This information was followed by a change or sustain talk statement, which participants listened to and could also read on the screen. Following the statement, participants made 15 food choices. Food items with varying levels of healthiness and tastiness were presented. Each choice trial started with food displayed on the screen, and participants had to rate on a 4-point Likert scale how much they wanted to eat the food for real at the end of the experiment. The choice was displayed on the screen when they hit a corresponding response button. A fixation cross-separated trials within each talk block. The duration of each event within these two example trials is displayed in seconds.

All food stimuli were selected from a database of 600 food pictures. The food pictures were presented on a computer screen in form of high-resolution pictures (72 dpi). MATLAB and Psychophysics Toolbox extensions (54) were used for stimulus presentation and response recording. Participants saw the food stimuli via a head-coil–based mirror and indicated their responses using an fMRI compatible response box system. The hand used to make choices and the display of the 4-point Likert scale (from left to right or right to left) was counterbalanced across participants. The task was incentive compatible, because one food was chosen out of chance for consumption at the end of the experiment (22, 31).

### Body composition and questionnaires

After the fMRI session, anthropometric measures such as body weight and body fat percentage were obtained via bio-impedance analysis using Tanita® Body Composition Analyzer SC-240MA (Tanita Corporation, Tokyo, Japan); height was self-reported. Body mass index (BMI) was computed as weight (kg)/height squared (m^2^) (Kg/m^2^). The habitual physical activity level was estimated via the short form of the International Physical Activity Questionnaire (IPAQ-SF), which assesses the frequency and duration of physical activity according to intensity categories (moderate, vigorous, and walking) and sitting time over the previous seven days. Estimates of total physical activity are given in metabolic equivalent task-min/week (MET-min/week) (55). Participants filled in the Yale Food Addiction Scale (YFAS) to measure signs of addictive-like eating behavior. It comprises 35 items with eight response options ranging from “Never” to “Every Day.” Each question assesses one of the 11 diagnosis criteria defined originally for substance abuse (e.g., tolerance, withdrawal, clinical Significance). The cut-off for food addiction is a total score of at least two of these 11 diagnosis criteria and meeting the criteria to be clinically significant (56).

### Dietary intake assessment

To ascertain the required nutrition-related variables, once the 24h-recall was collected, energy intake (in Kcal), sodium and fatty acids quantities were ascertained through Nutrilog SAS software (Nutrilog-Online 20231010). Additionally, to measure diet quality, the healthy eating index (HEI) was computed (57). The HEI-2015 is based on the Dietary Guidelines for Americans 2015-2020 (for a comparison between the American guidelines and national guidelines see *SI Appendix* Table S4). It has 13 items whose scoring follows adequacy- or moderation-based energy-adjusted intakes cut-offs, except for the fatty acids components. Each components weights differently into the final scoring (0-100, where higher scores indicate a healthier dietary pattern): total fruits (5 points), whole fruits (5 points), total vegetables (5 points), greens and beans (5 points), whole grains (10 points), dairy (10 points), total protein foods (5 points), fatty acids (polyunsaturated fatty acid plus monounsaturated fatty acid to saturated fatty acid ratio; (10 points), refined grains (10 points), sodium (10 points), added sugars (10 points), and saturated fats (10 points). Refined grains, sodium, added sugars, and saturated fats as components to be consumed in moderation have a reversed scoring (57). The HEI score for the sample is shown in Table 1.

### Imaging data acquisition

Brain images were collected on a 3 T Verio Siemens scan, using a 32-channel receive-only head coil. For the structural acquisition a whole-brain high-resolution T1-weighted structural scans were acquired for all participants with a magnetisation prepared rapid gradient echo (MPRAGE) sequence with the following parameters: TR = 2.4 s; TE = 2.05 ms; flip angle=9°, slice thickness=0.8mm. For the functional acquisition, a multi-band T2^*^-weighted multi-echoplanar images (mEPI) with BOLD contrast were acquired. To cover the whole brain with a repetition time of 1.25 seconds, the following sequence parameters were applied: echo times 14.8 ms; 33.4 ms and 52 ms; 48 slices collected in an interleaved manner; 3-mm slice thickness, FOV = 192 mm; flip angle = 68°, and an average of 646 volumes per participate session. Three echos were acquired to better accommodate for the compromise between spatial resolution and signal of the orbitofrontal context (58, 59). To further minimise the signal drop out in the orbitofrontal cortex, an oblique acquisition orientation of 30° above the anterior–posterior commissure line was applied (60).

### Imaging data pre-processing

Anatomical MPRAGE and functional mEPIs were processed using the Statistical Parametric Mapping – SPM12 – (Welcome Department of Imaging Neuroscience, Institute of Neurology, London, United Kingdom) using MATLAB version R2020b (The MathWorks Inc., United States). Each participant’s anatomical image was segmented into gray, white matter and cerebrospinal fluid using the SPM12 segmentation tool. The three echo images of each volume were summed into one EPI volume using the SPM 12 Image Calculator (61–63). Then, the summed EPIs were spatially realigned and motion corrected, coregistered to the mean image, normalised to the Montreal Neurological Institute (MNI) space using the same transformation as the anatomical image, and spatially smoothed using a Gaussian kernel with a full-width-at-half-maximum of 8 mm.

### Data analyses

Statistical tests were conducted with the Matlab Statistical Toolbox (Matlab 2020a, MathWorks), and JASP (JASP 0.16.1). Continuous variables were reported as mean and standard error of mean (SEM), and categorical variables as percentage.

### Behavioral analyses

Linear mixed-effects models (LMEs) were fit to mean-centered stimulus value (SV, coded -2 for “strong no”, -1 for “no”, 1 for “yes” and 2 for “strong yes”) ratings to test how much the type of talk (change talk coded 1, sustain talk coded -1) and the type of attribute (healthiness coded 1, tastiness coded -1) affected the valuation of food stimuli. Participant’s tastiness and healthiness ratings were used to categorize trials into healthy, tasty, unhealthy, and untasty choice trials. In total, two LMEs were fit to (1) healthy food items and tasty food items (LME 1), and (2) unhealthy food items and untasty food items trials (LME 2), respectively, and the following equation (I.):

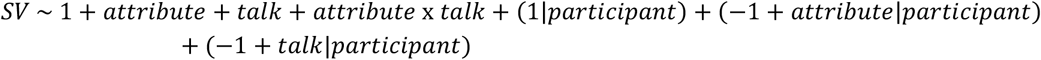

The model assumed random intercepts nested by participant number, and random slopes for the effect of type of talk (change versus sustain) and attribute (healthiness versus tastiness).

A similar third LME (LME 3) was fit to the neural craving signature responses to stimulus value (i.e., food choices) assigned to healthy and tasty food.

All LMEs were controlled for BMI. Post hoc, paired, one-tailed and two-tailed t-tests were used to determine the directionality of main effects and interactions.

To test how tastiness and healthiness attributes were integrated into stimulus value following change and sustain talk statements, the mean-centered stimulus values (SV, coded -2 for “strong no”, -1 for “no”, 1 for “yes” and 2 for “strong yes”), were fit to a multilevel general linear model (GLM) following equation II:

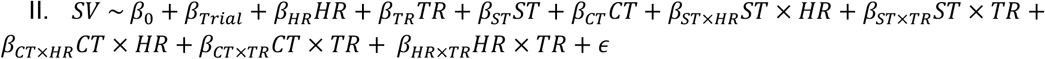

On the first level analyses, the model included the following regressors: trial number (β_Trial_) to account for fatigue effects, health ratings (β_HR_), and taste ratings (β_TR_), and two categorical indicators for sustain talk (β_ST_) and change talk runs (β_CT_). Additionally, the model included four interaction terms: STxHR, CTxHR, STxTR, CTxTR. These interactions tested for effects of healthiness and tastiness on stimulus value in trials following change and sustain talk, respectively. Individual beta coefficients were then fit into a second-level factorial analysis of variance (ANOVA) to test for main effects of talk (change versus sustain talk), attribute rating (healthiness versus tastiness rating), and the interaction talk type by attribute type. Post hoc, paired, two-tailed t-tests were used to determine the directionality of interactions.

### Imaging analyses

Analogous to the behavioral analyses, two multilevel general linear models (GLMs) were fit to fMRI time series.

To detect the voxels involved in SV encoding, a first GLM included on the first level four regressors of interest: (1) an indicator for type of talk with boxcar durations of 15 seconds, (2) choice onset regressor with a boxcar duration corresponding to reaction times, (3) a parametric modulator at time of choice consisting of stimulus value, and (4) missed trials with a boxcar duration of 3 seconds. The six realignment parameters were added as regressors of non-interest to correct for head motion. Individual beta images for this parametric modulator SV at the time of choice were then fit into a second-level random effects analysis using one-sampled t-tests.

To identify the voxels involved in the encoding of healthiness and tastiness as a function of the type of talk, a second GLM included nine regressors of interest: (1) indicators for onset of change talk statements with boxcar durations of 15 seconds, (2) choice onset regressors at time of food image display following change talk with durations corresponding to the reaction time, two parametric modulator regressors at time of food choice following change talk: (3) healthiness and (4) tastiness ratings, (5) indicators for onset of sustain talk with boxcar durations of 15 seconds, (6) food choice onset at time of food image display following sustain talk with a duration corresponding to the reaction time, and two parametric modulator regressors: (7) healthiness and (8) tastiness, (9) onset regressors for missed trials with boxcar duration of 3 seconds. The 6 realignment parameters were included as regressors of non-interest to control for head movement. Then, individual beta images were fit into a second-level random effects analysis using one-sampled t-tests, adding BMI as a covariate to the matrix.

### Definition of regions of interest (ROI) for small volume correction

Regions of interest for small volume correction were functionally defined by the significant, whole brain family wise error corrected activation within the vmPFC by stimulus value across all trials (MNI = [-6, 50, -8], *SI Appendix* Table S2).

### Responses of the neurobiological craving signature (NCS) to food stimulus value (i.e., choices)

To assess how change and sustain talk altered craving-related brain responses (as measured by the NCS, Koban et al., 2023), a general linear model (GLM 3) fit each participant’s BOLD signal. It included ten regressors of interest: (1) an indicator for type of talk, (2) an onset regressor for tasty food choices following change talk, (3) stimulus value as a parametric modulator of the tasty food choice onset regressor following change talk, (4) an onset regressor for healthy food choices following change talk, (5) stimulus value as a parametric modulator of the healthy food choice onset regressor following change talk, (6) an onset regressor for tasty food choices following sustain talk, (7) stimulus value as a parametric modulator of the tasty food choice onset regressor following sustain talk, (8) an onset regressor for healthy food choices following sustain talk, (9) stimulus value as a parametric modulator of the healthy food choice onset regressor following sustain talk, and onset regressors for missed trials with boxcar duration of 3 seconds. The six realignment parameters were included as regressors of non-interest to control for head movement.

NCS responses were quantified as the dot product between the contrast images of interest (i.e., on the parametric regressors 3 and 5 for tasty and healthy stimulus value at time of food choice) for each participant and the NCS weight map, plus a fixed intercept. This results in one scalar value per contrast image and participant, which can be further analyzed using standard statistical tests.

## Data availability

Behavioral data is available on the study’s OSF website.

## Code availability

Code to analyse behavioral data and food decision making task scripts are available on study’s OSF website. The NCS and code to apply it to any fMRI data (apply_ncs.m function) are publicly available on Github (https://github.com/canlab/Neuroimaging_Pattern_Masks/tree/master/Multivariate_signature_patterns/2022_Koban_NCS_Craving).

## Acknowledgments

We thank Hilke Plassmann and Nathalie George for their insightful project discussion. We thank the CENIR team for their support and collaboration throughout the project, Valentine Lemoine (clinical research assistant, ICAN) and the platform ICAN for help in clinical investigation (recruitment). This project has received funding from the European Union’s Horizon 2020 research and innovation program under the Marie Skłodowska-Curie grant agreement n° 945298. BR is a Fellow of the Paris Region Fellowship Programme supported by the Paris Region. LK acknowledges funding from an ERC Starting Grant (SOCIALCRAVING, 101041087). However, the views and opinions expressed are those of the authors only and do not necessarily reflect those of the European Union or the European Research Council. Neither the European Union nor the granting authority can be held responsible. The funders had no role in study design, data analysis, manuscript preparation, or publication decisions. LS acknowledges funding from Fondation NRJ – Institute de France.

