## Supporting Information for "How motivational interviewing shifts food choices and craving-related brain responses to healthier options"

[e] Departement de Psychiatrie Adulte, Hôpital Pitié-Salpêtrière, Assistance Publique Hopitaux Publiques (AP-HP).

[f] Centre de Recherche en Neurosciences de Lyon (CRNL), CNRS, INSERM, Université Claude Bernard Lyon 1, Bron, France

* Belina Rodrigues and Liane Schmidt

**Table S1.** Multiple regression results in stimulus value (SV) for all participants.

| N=32 | Intercept | Trial | HR | TR | ST | CT | ST*HR | ST*TR | CT*HR | CT*TR | HR*TR |
|---|---|---|---|---|---|---|---|---|---|---|---|
| **Coeff** | -0.28 | 0.00 | 0.12 | 0.32 | -0.29 | -0.29 | -0.01 | 0.22 | 0.16 | 0.13 | 0.02 |
| **STE** | 0.08 | 0.00 | 0.02 | 0.02 | 0.05 | 0.05 | 0.02 | 0.02 | 0.03 | 0.02 | 0.01 |
| **T** | -3.54 | 0.17 | 6.20 | 17.94 | -6.31 | -6.27 | -0.42 | 10.98 | 5.72 | 6.62 | 1.39 |
| **Z** | -3.21 | 1.13 | 3.66 | 4.18 | -3.34 | -2.91 | -0.91 | 4.59 | 3.99 | 3.19 | 1.02 |
| **P** | 0.0007 | 0.802 | 0.0001 | 0.0000 | 0.0004 | 0.0018 | 0.8195 | 0.0000 | 0.0000 | 0.0007 | 0.1544 |

**Table S2.** Regions positively correlated with stimulus value independent of the condition (Whole-brain FWE<0.05).

| Region | BA | K | X | Y | Z | Peak Z score |
|---|---|---|---|---|---|---|
| dACC | 32 | 9 | -2 | 28 | -10 | 5.07 |
| **vmPFC** | 10 | **38** | **-6** | **50** | **-8** | **4.93*** |
| | | 8 | -2 | 70 | 6 | 4.81 |
| Ventral Posterior Cingulate | 23 | 17 | -2 | -58 | 16 | 4.80 |

* and highlight in bold corresponds to the MNI coordinates used for small volume correction.

**Table S3**. Example de phrases.

| Example of sentences of a participant who wanted to decrease the amount of chips |
|---|
| **Sustain talk:** "I do not see an immediate effect. So, I tell myself " It's is not that bad, I will finish this bowl of chips". |
| **Change talk:** "In terms of weight, I find that I have put on some weight recently, and I think it has to be that." |

**Table S4**. Comparison between HEI-2015 scoring, based on the Dietary guidelines for Americans 2005-2010 vs national dietary recommendations.

| | COMPONENT | STANDARD FOR MAXIMUM SCORE | STANDARD FOR MINIMUM SCORE | RECOMMENDATIONS |
|---|---|---|---|---|
| ADEQUACY | **Total Fruits[1]** | ≥0.8 c.eq[2]./1,000 kcal | No Fruits | - At least 5 fruit and vegetables per day;<br>- A small handful of nuts each day: unsalted walnuts, hazelnuts, almonds, pistachios, etc; |
| | **Whole Fruits** | ≥0.4 c.eq./1,000 kcal | No Whole Fruits | |
| | **Total Vegetables[3]** | ≥1.1 c.eq./1,000 kcal | No Vegetables | |
| | **Greens and Beans[3]** | ≥0.2 c.eq. per 1,000 kcal | No Dark Green Vegetables or Legumes | - Pulses at least twice a week; |
| | **Whole Grains** | ≥1.5 c.eq. per 1,000 kcal | No Whole Grains | - At least one wholegrain starch per day; |
| | **Dairy[4]** | ≥1.3 c.eq. per 1,000 kcal | No Dairy | - 2 dairy products per day; |
| | **Total Protein Foods[3]** | ≥2.5 oz eq. per 1,000 kcal | No Protein Foods | - Favour poultry and limit other meats (pork, beef, veal, mutton, lamb, offal) to 500g per week;<br>- Limit charcuterie to 150g per week<br>- Fish twice a week, including fatty fish (sardines, mackerel, herring, salmon);<br>-A small handful of nuts each day: unsalted walnuts, hazelnuts, almonds, pistachios, etc; |
| | **Seafood and Plant Proteins[5]** | ≥0.8 oz eq. per 1,000 kcal | No Seafood or Plant Proteins | - Fish twice a week, including fatty fish (sardines, mackerel, herring, salmon);<br>- A small handful of nuts each day: unsalted walnuts, hazelnuts, almonds, pistachios, etc. |
| | **Fatty Acids** | (PUFAs[6] + MUFAs7)/SFAs[8] ≥2.5 | (PUFAs + MUFAs)/SFAs ≤1.2 | - Added fats - oil, butter and margarine - can be consumed every day in small quantities. Favour rapeseed, nut and olive oil; |
| MODERATION | **Refined Grains** | ≤1.8 oz eq. per 1,000 kcal | ≥4.3 oz equiv. per 1,000 kcal | - At least one wholegrain starch per day; |
| | **Sodium** | ≤1.1 gram per 1,000 kcal | ≥2.0 grams per 1,000 kcal | - Reduce salt consumption; |
| | **Added Sugars** | ≤6.5% of energy | ≥26% of energy | - Limit sugary drinks, fatty, sugary, salty and ultra-processed foods; |
| | **Saturated Fats** | ≤8% of energy | ≥16% of energy | - Added fats - oil, butter and margarine - can be consumed every day in small quantities. Favour rapeseed, nut and olive oil; |

**1:** Includes 100% fruit juice; **2 -** c.eq - cup equivalent; **3:** Includes legumes (beans and peas); **4:** Includes fortified soy beverages.; **5:** Includes seafood, nuts, seeds, soy products (other than beverages), and legumes (beans and peas).; **6**: polyunsaturated fatty acids; **7**: monounsaturated fatty acids; **8:** saturated fatty acids.
